# Effects of sequence-dependent non-native interactions in equilibrium and kinetic folding properties of knotted proteins

**DOI:** 10.1101/2023.06.06.543861

**Authors:** João N. C. Especial, Patrícia F. N. Faísca

## Abstract

Determining the role of non-native interactions in folding dynamics, kinetics and mechanisms is a classical problem in protein folding. More recently, this question has witnessed a renewed interest in light of the hypothesis that knotted proteins require the assistance of non-native interactions to fold efficiently. Here, we conducted extensive equilibrium and kinetic Monte Carlo simulations of a simple off-lattice C-alpha model to explore the role of non-native interactions in the thermodynamics and kinetics of three proteins embedding a trefoil knot in their native structure. We find that equilibrium knotted conformations are stabilized by non-native interactions that are non-local, and proximal to native ones, thus enhancing them. Additionally, non-native interactions increase the knotting frequency at high temperature, and in partially folded conformations below the transition temperature. While non-native interactions clearly enhance the efficiency of the transition from an unfolded conformation to a partially folded knotted one, they are not required to efficiently fold a knotted protein. Indeed, a native centric interaction potential drives the most efficient folding transition, provided that the simulation temperature is well below the transition temperature of the considered model system.

## 1 Introduction

Knotted proteins are globular proteins that embed a physical (i.e. open) knot in their native structure. The first knotted protein was identified in 1977 [1], and according to the most recent survey, ≈ 1% of the protein entries deposited in the Protein Data Bank (PDB) correspond to knotted chains [2]. More recently, a study that investigated protein structures predicted with AlphaFold [3] found that knots are present in 0.2% of the human proteome [4]. Knotted proteins turn out with different levels of topological complexity, from the simplest and most abundant trefoil (or 3_1_) knot, to the remarkably complex 6_3_ knot, exclusively found in a human protein [5]. Protein knots are further classified as deep or shallow, according to whether the number of residues that must be removed from one of the termini to unknot the chain is larger or smaller than 10, respectively [6].

Despite being statistically rare [7], knotted proteins have attracted considerable attention, and during the last 20 years a plethora of experimental and theoretical investigations has been dedicated to understanding how they knot and fold (reviewed in [6, 8]), and to establish their biological role (reviewed in [9]).

The very first computational study addressing knotted proteins [10], investigated the folding and knotting mechanism of protein YibK, which features a deep trefoil knot. The study used a C-alpha model combined with a native-centric Gō potential, and sampling was performed with fixed temperature Langevin Molecular Dynamics (MD) simulations. Interestingly, and quite surprisingly, it was found that in timescales typically used in molecular simulations, no knotting event was observed in any of the considered trajectories, suggesting that native interactions are not sufficient to knot the protein in a reasonable timescale. Another study focusing on the same protein [11], and using the same model and sampling methodology, reported an exceedingly small (1-2%) knotting frequency in the attempted MD trajectories. However, a modified native centric Gō potential in which non-native interactions were applied selectively to two chain segments (86-93 and 122-147) of YibK, was found to fold 100% of the attempted trajectories [10]. These non-native interactions were identified by trial and error. A subsequent study framed on fixed temperature Monte Carlo (MC) simulations of a C-alpha model, explored AOTCase, a protein larger than YibK, which also embeds a deep trefoil knot in its native structure [12]. Again, no knotting event was recorded when protein folding energetics was modelled through a native-centric Gō potential, but a modified Gō potential with sequence-dependent non-native interactions (i.e., non-native interactions that establish between any pair of contacting residues and not only between selected residues as in [10]) was found to produce a detectable fraction of knotted structures in partially folded conformations (when only ≈ 30% of the native contacts are formed).

The situation is different for shallow knotted proteins on-lattice. In this case a standard Gō potential efficiently folds model systems featuring shallow knots (including a 5_2_ knot) in the context of MC simulations conducted at fixed temperature, although knotted proteins require far more MC steps to fold than their unknotted counterparts [13, 14, 15]. Moreover, off-lattice MC simulations of MJ0366, the smallest knotted protein found to date featuring a shallow trefoil knot, were also successful in correctly folding the chain, both at fixed temperature and with a replica-exchange engine [16, 17].

The results summarized above thus seem to indicate that non-native interactions are necessary to fold proteins featuring deep knots in a biological timescale, but how they succeed in doing so is not clear. As a matter of fact, understanding the role of non-native interactions in the folding of knotted proteins has been pointed out as a major fundamental question by some authors in recent years [8].

For several proteins knotting proceeds through a single event in which one of the termini threads through a loop, the so-called knotting loop, formed by the remainder of the chain. It is possible that non-native interactions serve to enhance the native interactions that already exist between the regions involved in knotting as suggested for YibK [10], that they stabilize the transition state as shown by a combination of experiments and simulations on MJ0366 [18], that they may assist threading events, or stabilize knotted intermediate states as suggested by lattice models [19].

Here we explore the role of native and non-native interactions in both equilibrium and kinetic properties of the folding transition of three knotted trefoil proteins by conducting extensive Monte Carlo simulations of a simple C-alpha off-lattice model. We consider two interaction potentials: a purely native-centric Gō potential, and a modified Gō potential in which sequence-dependent non-native interactions are added as perturbation to the stronger native ones. Equivalent strategies have been used elsewhere [20, 21, 22]. We found that non-native interactions can drive a downhill folding transition. Additionally, non-native interactions can substantially increase the knotting frequency above the transition temperature, and in partially folded conformations below the transition temperature. Equilibrium knotted conformations are stabilized by non-native interactions that are non-local, and proximal to native ones, thus enhancing them. Kinetic simulations show that the pure Gō potential is able to efficiently fold knotted proteins, provided the simulation temperature is well below the transition temperature. However, in that temperature range, non-native interactions drastically increase the efficiency of the transition from unfolded to a partially folded knotted conformation.

## 2 Models and methods

### 2.1 Protein representation

To represent the protein conformation we use a simple C-alpha model in which residues are coarsegrained as hard spherical beads of uniform size, centered on the C-alpha atoms. These, in turn, constitute spherical joints that articulate rigid sticks which represent the amide planes that connect consecutive C-alpha atoms in the backbone. We adopt a radius of 1.7 ÅA for the beads, which is the van der Waals radius of C-alpha atoms [23], and for the length of each stick we adopt the distance between the C-alpha atoms of the respective bonded residues in the protein ‘s native conformation, these being approximately 2.9 ÅA, for cis bonds, and 3.8 ÅA, for trans bonds. Two non-bonded residues are said to be in contact in the native conformation if the smallest distance between any two heavy atoms, one belonging to each residue, is ⩽ 4.5 ÅA, this cut-off being chosen because it is slightly larger than twice the average van der Waals radius of heavy atoms in proteins.

### 2.2 Interaction potential

#### 2.2.1 Gō potential

For a purely native-centric Gō potential, the total energy of a conformation defined by bead coordinates 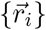 is given by

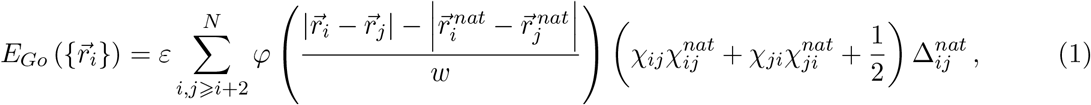

where 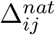 is the native contact map matrix, that takes the value 1 if the *i* − *j* contact is present in the native conformation and is 0 otherwise, *ε* is a uniform intramolecular energy parameter (taken as *−*1 in this study, in which energies and temperatures are shown in reduced units), N is the chain length measured in number of beads, *φ* is the potential well associated with the native contacts, w is the half-width of this potential well, and the chirality of contact *i* − *j* in the conformation under consideration is

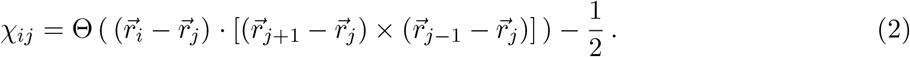

The chirality of the *i* − *j* contact in the native conformation is

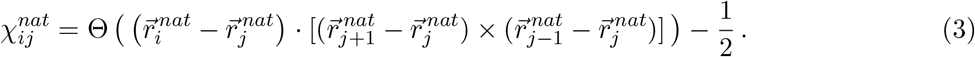

In equations (2) and (3), Θ is Heaviside ‘s unit step function, which takes the value 1 if its argument is greater than zero and the value 0 otherwise. The chirality factor in (1) favors the native conformation *vis a vis* its mirror conformation, thereby ensuring chirality coherence among all contacts and convergence of the simulations towards the native ensemble for temperatures below transition temperature.

In this study we use inverse quadratic potential wells and equation (1) becomes

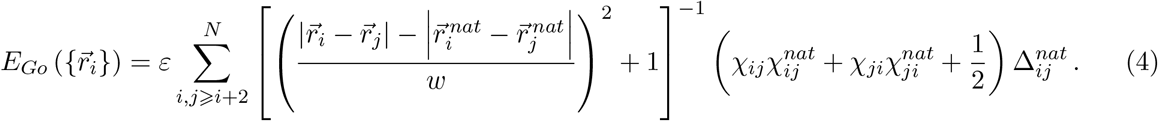

The half-width of the potential well, *w*, determines the degree of cooperativity of the folding transition, with a wider well leading to less cooperative transitions occurring at higher transition temperatures. We measure the degree of cooperativity of the transition by the ratio of the full width at half maximum (FW HM) of the *C*_*V*_ peak to the temperature at which the peak occurs, the melting temperature, *T*_*m*_. A typical two-state transition that has been well characterized experimentally is that of the B1 domain of protein G (PDB id: 2GB1 [24]) and its *FW HM/T*_*m*_ ratio at pH 5.4 has been determined to be approximately 4.4% [25]. Hence, the half-width of the potential well is adjusted to obtain a simulated *FW HM/T*_*m*_ ratio between 4 and 5%. This is not just a technical detail of the model since very narrow transitions, which lead to artificially large cooperativity, are poorly sampled in equilibrium simulations that use a replica exchange procedure as done here [16, 26, 27]. The width proposed above has been successfully used in previous simulations employing a similar potential and sampling [16, 17, 28, 29]. When this potential is used, we consider that a native contact is formed if the distance between the centers of the respective beads differs from the distance between their C-alpha atoms in the native conformation by less than the half-width of the potential wells, w.

#### 2.2.2 Modified Gō potential

The modified Gō potential is a linear combination of the Gō potential described above with a sequence-dependent (SD) potential term, 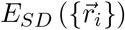, that accounts for non-native interactions. In the modified Gō potential the total energy of a conformation 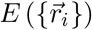 defined by bead coordinates 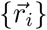 is given by

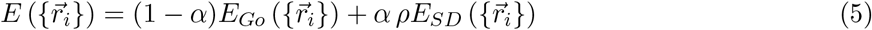

with

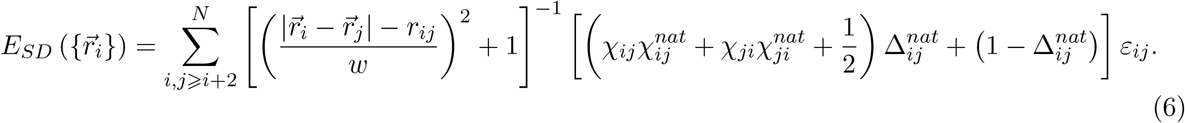

In the equation above *ε*_*ij*_ is an interaction parameter calculated through a least-squares fit to the native contact matrix (see Appendix). The least-squares fit ensures that this potential is the least disruptive SD potential that may be added as perturbation to the Gō potential. If *i* − *j* is a native contact, 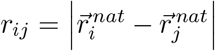 and, if *i* − *j* is a non-native contact, *r*_*ij*_ is approximated by the arithmetic mean of all native contact distances. To identify which native and non-native contacts are formed in a sampled conformation we consider that contact *i* − *j* is formed if the distance between the centers of the beads *i* and *j* differs from *r*_*ij*_ by less than the half-width of the potential wells, w.

In equation (5) the parameter 0 ⩽ *α* ⩽ 1 controls the strength of the non-native interactions and the constant *ρ*, defined as

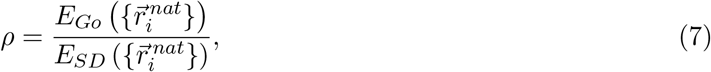

normalizes the SD potential term to the same native energy as the Gō potential, thus ensuring that native energy remains unchanged upon adding this perturbative term, whatever the value of *α*.

### 2.3 Monte Carlo sampling

The conformational space is explored with Metropolis [30] Monte Carlo method by using a move set that comprises two elementary moves, crankshaft (Figure 1A) and pivot (Figure 1B).

**Figure 1:**
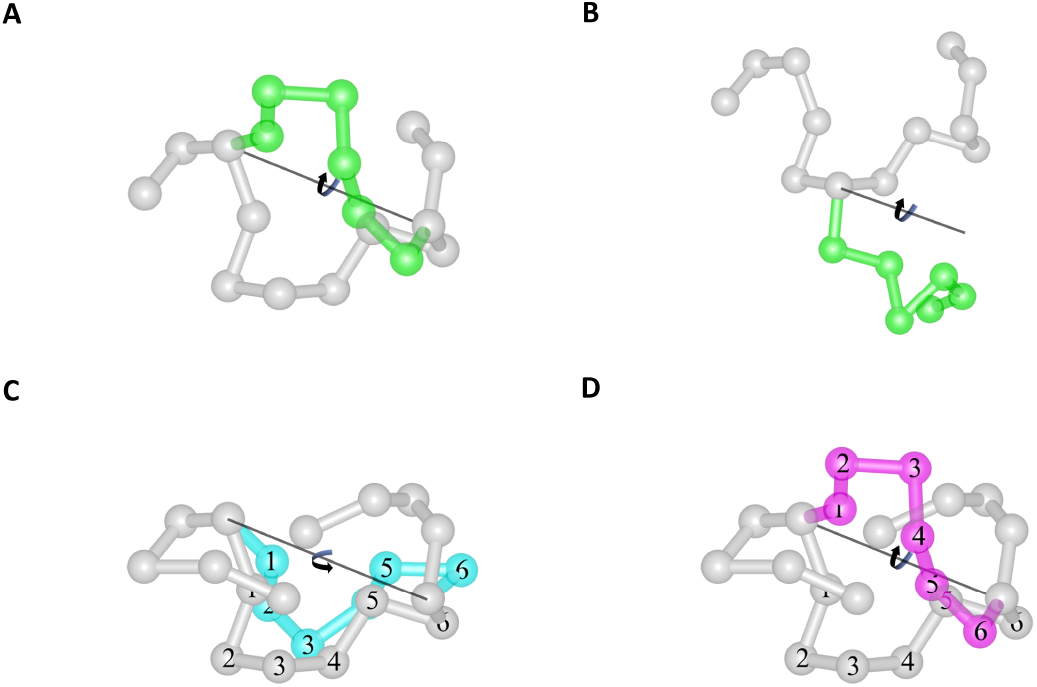
The move set. In the crankshaft move (A) an axis passes through two randomly selected beads and the beads between these are rotated around the axis. In the pivot move (B) an axis passes through one randomly selected bead and the beads between that and a randomly selected terminus are rotated around the axis. Both moves have two variants: In the LTyP move variant (C) the rotated segment reaches the cyan conformation without passing across any fixed segment, hence preserving the linear topology of the chain throughout the move; In the non-LTyP move variant (D) the rotated segment can only reach the magenta conformation by either moving across the front or across the back fixed segments and, hence, the final conformation, free of steric clashes, requires that the linear topology of the chain not be preserved throughout the move.

These elementary moves can be performed in either of two ways: 1) by limiting the amplitude of the rotation so that no bead or stick is allowed to overlap or move across another (e.g. Figure 1C); 2) by not limiting the rotation and allowing such crossings to potentially occur (e.g. Figure 1D) as long as no steric clashes are present in the resulting trial conformation. A simulation that only performs moves of the first kind preserves the linear topology of the chain in all Monte Carlo steps and is designated as a linear topology preserving (LTyP) simulation. Conversely, a simulation that allows moves that take the chain across itself is designated as non-LTyP.

#### 2.3.1 Equilibrium simulations

To sample canonically distributed conformational states we use Metropolis [30] Monte Carlo simulations combined with replica-exchange (or parallel tempering) [31, 32] (MC-RE). For complex structures, such as those of deeply knotted proteins, the preservation of linear topology provides correct equilibrium results but only after long relaxation. Indeed, as shown in [33] they require up to two orders of magnitude more Monte Carlo steps to equilibrate than non-topology preserving simulations of the same model system. Thus, in the case of equilibrium simulations, we use the non-LTyP variant of the move set. Further details on the RE-MC simulations can be found elsewhere [33].

The Weighted Histogram Analysis Method (WHAM) [34] was used to analyze data generated by the MC-RE simulations and produce maximum likelihood estimates of the density of states, from which expected values for thermodynamic properties were calculated as functions of temperature. In particular, the heat capacity, *C*_*V*_, defined in reduced units as C_*V*_ = (< *E*^2^ > − < *E* >^2^)/*T* ^2^. The melting temperature, *T*_*m*_, was determined as the temperature at which the *C*_*V*_ peaks and the width of this peak at half-maximum was determined to calculate the *FW HM/T*_*m*_ ratio.

The fraction of native contacts, *Q*, was chosen as the reaction coordinate and WHAM was also used to project the density of states along *Q* to obtain free energy and knotting probability profiles along this coordinate at a specific temperature.

In both equilibrium and kinetic simulations (see below) the topological state (knotted or unknotted) of a sampled conformation was determined using the Koniaris-Muthukumar-Taylor (KMT) algorithm [35]. The temperature at which p_*K*_ = 0.5 is regarded as the central temperature of the knotting transition and designated as T_*k*_.

#### 2.3.2 Kinetic simulations

The physical process of protein folding is subject to excluded volume. Therefore, in order to properly capture the sequence of conformational transitions that establish a pathway starting from an initial unfolded conformation to a final knotted conformation, it is necessary to use the LTyP variant of the move set. Thus, kinetic simulations are performed with the LTyP variant of the move set at fixed temperature.

In this work we are interested in assessing folding efficiency and knotting efficiency as a function of temperature. In both cases, the MC simulation starts from an unfolded conformation that was obtained by performing a MC simulation at high temperature (*T* = 4.0). Such conformation is unknotted and its *Q* is smaller than the *Q* at which the knotting probability (at *T*_*k*_ and for the Gō potential) is 0.01. When the purpose of the simulation is to access folding efficiency it stops when the native conformation is found. The criteria used to determine if a conformation is native is quite stringent: a conformation is deemed native if it is knotted and its *Q* is larger than that at which the knotting probability (at *T*_*k*_ and for the Gō potential) is 0.99.

To investigate how folding and knotting efficiency depend on temperature we conducted kinetic simulations at several temperatures. For all considered model systems 24 kinetic simulations were performed at temperatures below *T*_*m*_, and for proteins MJ0366 and Rds3p, 8 additional simulations were performed at *T*_*m*_ and temperatures above. All temperatures were separated by constant increments equal to the *FW HM* of the *C*_*V*_ peak. Since the number of MC steps does not correspond to a physical time, instead of considering folding time (or, in alternative, folding rate) we calculated the number of unfolding to folding (or knotting) transitions that occurred per million mcs (Mmcs) at each considered temperature. A model system that folds (or knots) more efficiently will exhibit a higher number of transitions at a certain temperature.

## 3 Results

### 3.1 Model systems

We consider three proteins featuring a trefoil knot in their native structures.

Protein MJ0366 from *Methanocaldococcus jannaschii* (PDB id: 2efv) [36] is the smallest knotted protein found to date. It has 92 residues, and the knotted core comprises residues 11 to 82. Since both knot tails are short (10 residues), the knot is classified as shallow. The shallow knot is easily perceived in the native structure, being formed by threading the C-terminal helix, alpha-4, through the knotting loop formed by helices alpha-1 and alpha-2 (Figure 2A). The second protein is Rds3p (PDB id: 2k0a) [37], a metal binding protein from *Saccharomyces cerevisiae*. Rds3p is 109 residues long and the knotted core extends from residue 21 to residue 74. The N-tail of the knot is 20 residues long, and the C-tail comprises 35 residues (Figure 2B). Therefore, the knot is classified as deep. The third protein is YibK (PDB id: 1j85) [38], a methyltransferase from *Haemophilus influenzae*. This is the largest protein investigated in this study, featuring 160 residues with the knotted core extending from residue 77 to residue 119. In this case the N-tail is 76 residues long and the C-tail 41 residues long (Figure 2C). The knot classifies as deep.

**Figure 2:**
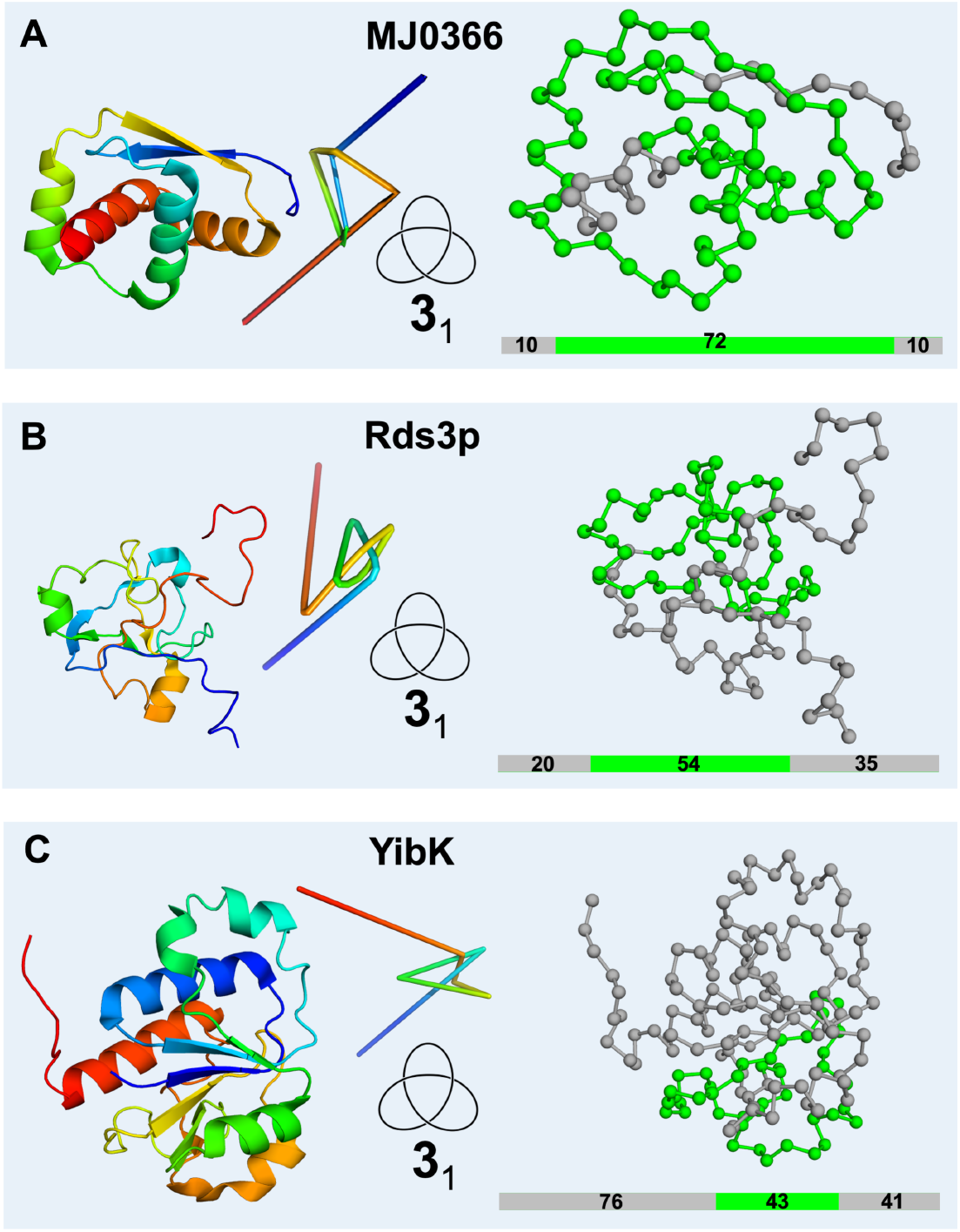
Model systems used in the present study. Cartoon representation (left) and bead and stick representation (right) of the native structure of proteins MJ0366 (A), Rds3p (B), and YibK (C). Also shown is the reduced knot representation and a diagrammatic representation of the polypeptide chain highlighting the size of the knotted core (in green) and that of the knot tails (in grey).

### 3.2 Non-native interactions and the nature of the folding transition

The effects of non-native interactions in decreasing the free energy barrier associated with a two-state folding transition have been studied theoretically [20] and observed in molecular simulations of several model systems [20]. In particular, a downhill scenario is predicted by the free energy landscape theory when the entropic and enthalpic contributions to the free energy compensate each other to the point where no significant (> 3*RT*) barrier appears along the folding reaction coordinate [39, 40, 41].

In Figure 3 we report the temperature dependence of the heat capacity, *C*_*V*_, energy, *E*, and knotting probability, *p*_*K*_, for proteins MJ0366 (left), Rds3p (center) and YibK (right) for the pure Gō and modified Gō potentials with increasing strength (*α* = 0.05, 0.10, 0.15 and 0.20) of non-native interactions. Additionally, we also show the projection of the free energy on the reaction coordinate *Q* at the temperature *T*_*k*_. This temperature is nearly identical to *T*_*m*_ except for MJ0366 with *α* = 0.20 due to the higher peak that occurs in its *C*_*V*_ at much lower temperature. For the three considered model systems the adopted parametrization of the pure Gō potential delivers a two-state folding transition. We observe the flattening of the heat capacity curves and the progressive lowering of the free-energy barrier as the strength of non-native interactions increases such that for *α* = 0.15 and *α* = 0.10 the transition becomes effectively downhill for MJ0366 and YibK, respectively. In the case of MJ0366 the heat capacity curve exhibits a low temperature peak that corresponds to a structural rearrangement whereby the protein in a near-native conformation gets rid of residual non-native interactions. Interestingly, for protein Rds3p the two-state transition is conserved, despite the significant decrease of the free energy barrier. Consistently, the dependence of E on T conserves its sigmoidal shape in the case of Rds3p, but it becomes progressively less sigmoidal for the other model systems as non-native interactions become more prominent. The dependence of E on T also shows that non-native interactions contribute to energetically stabilize the partially folded (or denatured) conformations sampled at high temperature above the transition midpoint, which explains why the temperature at which the heat capacity peaks increases through the addition of non-native interactions.

**Figure 3:**
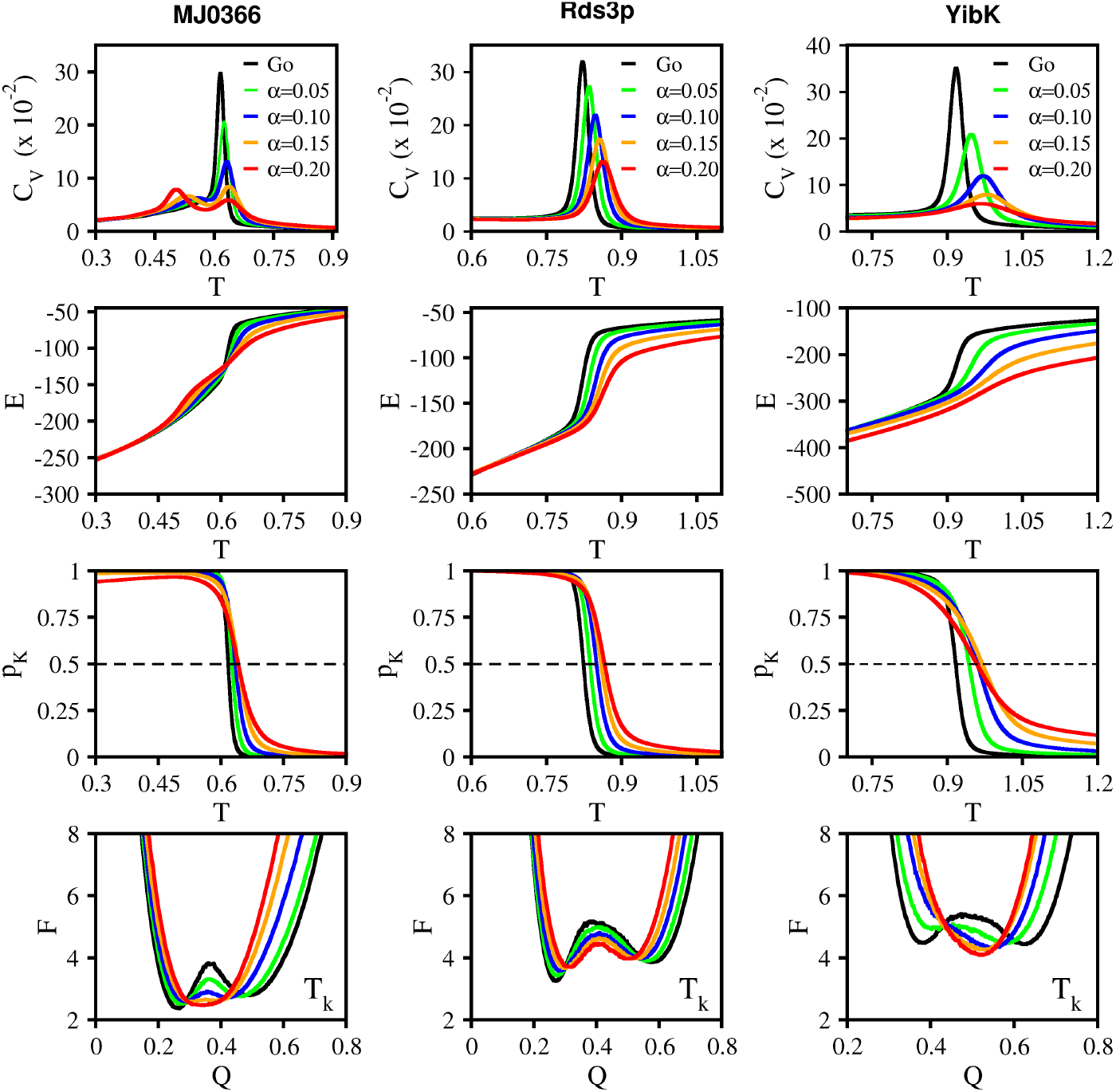
Folding transition. Dependence of the heat capacity, *C*_*V*_, energy, *E*, and knotting probability, *p*_*K*_, on temperature *T*, for proteins MJ0366 (left), Rds3p (middle) and YibK (right). The bottom row shows the projection of the free energy, *F*, on the reaction coordinate *Q* at temperature *T*_*k*_, which is the temperature at which *p*_*k*_ = 0.5.

It is also interesting to note that in the free energy profiles, the free energy minima corresponding to the thermally denatured ensemble shifts to the right as the strength of non-native interactions increases; this means that the thermally denatured ensemble becomes progressively more structurally consolidated through the addition of non-native interactions. At *T*_*k*_ the downhill transition relaxes to an equilibrium ensemble with approximately half of the native interactions established but the two-state transitions exhibit a more structurally consolidated native ensemble.

### 3.3 Non-native interactions and knotting probability

The analysis of the knotting probability curves in Figure 3 shows that their sigmoidal shape is fairly conserved across the considered interaction potentials (i.e. considered *α* values). However, the probability *p*_*K*_ to find knotted conformations at high temperatures (*T* > *T*_*k*_) increases significantly as the strength of non-native interactions is also increased (Figure 3). This behavior is consistent with the energetic stabilization driven by non-native interactions at high temperature, which compensate the entropic cost of knotting thus making it more favorable in that temperature regime.

We have also analyzed the dependence of *p*_*K*_ on *Q* at temperature *T*_*k*_ (Figure 4). We designate by *Q*_*k*_ the reaction coordinate for which knotting probability is 0.5. The fraction of native contacts *Q*_*k*_ varies between 0.35 (for MJ0366 with potential *α* = 0.20) and 0.54 (for YibK with Gō potential). Interestingly, when the folding transition becomes downhill the probability to find knotted conformations increases significantly below the transition midpoint (i.e., for *Q* < *Q*_*k*_), for proteins MJ0366 and YibK, presumably because non-native interactions substitute native ones, enabling stable knotted conformations to form with a smaller number of native interactions.

**Figure 4:**
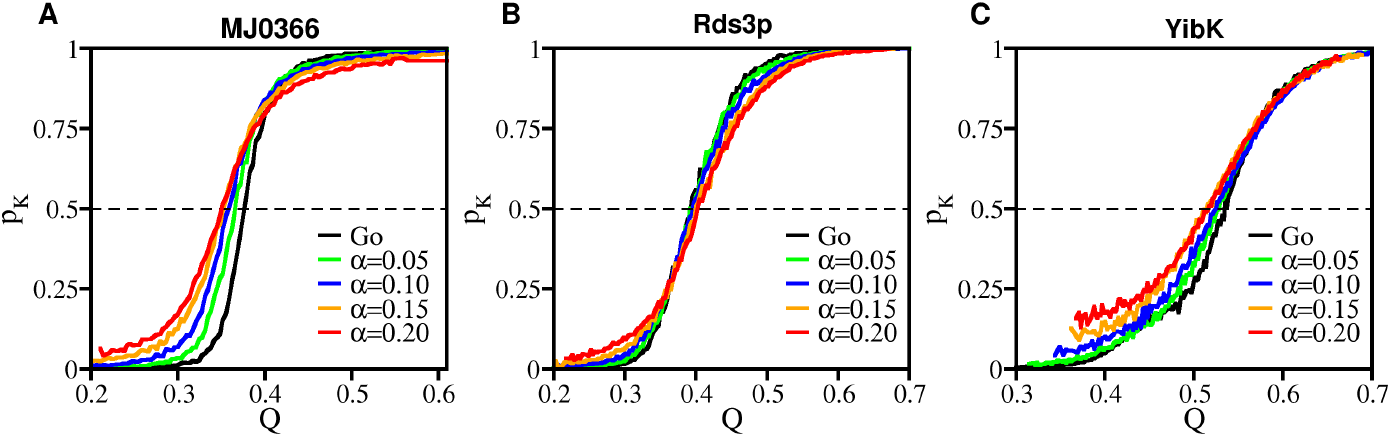
Knotting probability. Dependence of the knotting probability on the fraction of native contacts at *T*_*k*_. The reaction coordinate for which knotting probability is 0.5 is designated as *Q*_*k*_.

### 3.4 Non-native interactions that stabilize the knot

A conditional probability contact map is an *N* × *N* matrix whose elements represent the probability of any contact being formed in an equilibrium ensemble of conformations with the same topological state (knotted or unknotted) and fraction of native contacts *Q* at temperature *T*. To identify the native and non-native contacts that most likely contribute to the stabilization of knotted conformations we considered ensembles of conformations with *Q*_*k*_ at *T*_*k*_. By subtracting the conditional contact map of unknotted conformations from that of knotted ones and taking only the positive values, a differential contact map is calculated that can be used to identify the native and non-native interactions having the highest differential probability. The latter, which are reported in Figure 5, are the ones that most likely contribute to knot stabilization.

**Figure 5:**
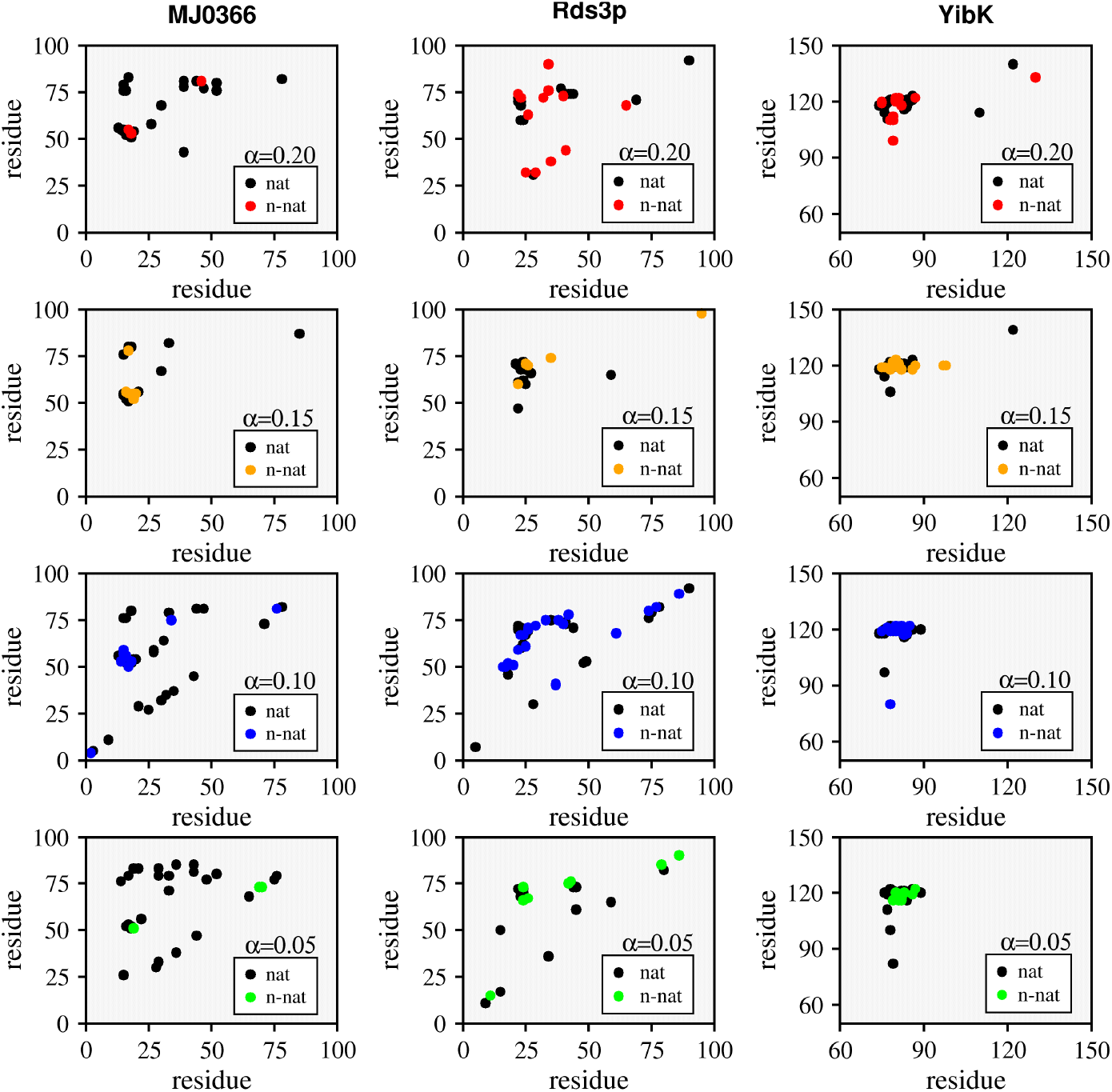
Interactions that stabilize the knot. Contact maps showing the native (black dots) and non-native (colored dots) contacts that appear with highest differential probability in the ensemble of knotted conformations with *Q*_*k*_ at *T*_*k*_ for proteins MJ0366 (left), Rds3p (middle) and YibK (right) for several strengths (*α* = 0.05, 0.10, 0.15 and 0.20) of non-native interactions

.The analysis of Figure 5 shows that the number of non-native interactions that stabilize knotted conformations is higher for the two proteins with deep knots. This is in line with previous observations that indicated that proteins with shallow knots do not require the assistance of non-native interactions to efficiently tie their backbones. Additionally, as the strength of non-native interactions increases the number of native local interactions (located close to the main diagonal) that stabilize the knotted conformations decreases. This is consistent with the fact that knotting requires interactions between residues that are located far away from each other in the polypeptide chain. The observed non-native interactions are thus mainly non-local being in tandem, or located nearby, native ones. This suggests that non-native interactions may operate as a system of redundant interactions that assist knotting, e.g., by keeping the protein in a conformation favorable for threading to occur. For the three proteins it is possible to identify clusters of native and neighboring non-native interactions that are broadly conserved across different *α*s (i.e. for different non-native interaction strength). For YibK these non-native interactions establish between residues located on the two stretches (75-87) and (119-122) of the polypeptide chain. In the case of protein MJ0366 the equivalent interactions involve residues located on the stretches (17-18) and (53-56). The interactions that stabilize the knot in the case of Rds3p appear to be more plastic than in the other two proteins although there is some level of conservation for the regions (22-34) and (63-76). Figure 6 highlights these regions in the three-dimensional native structures of the three proteins.

**Figure 6:**
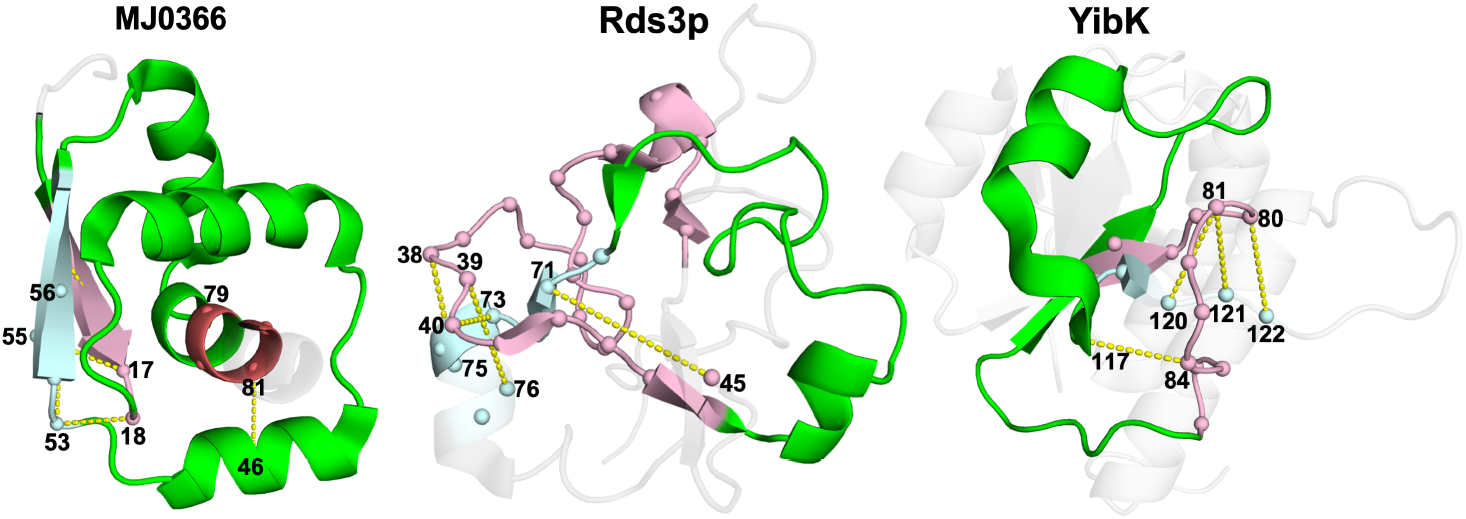
Interactions that stabilize the knot. Three-dimensional structures of MJ0366 (A), Rds3p (B) and YibK (C). In the cartoon representation only the knotted core is colored. Different colors are used to identify different chain segments of the knotted core containing residues involved in the most frequent native and non-native interactions that stabilize knotted conformations. Selected non-native interactions are represented as dotted yellow lines in the three-dimensional representation.

**Figure 7:**
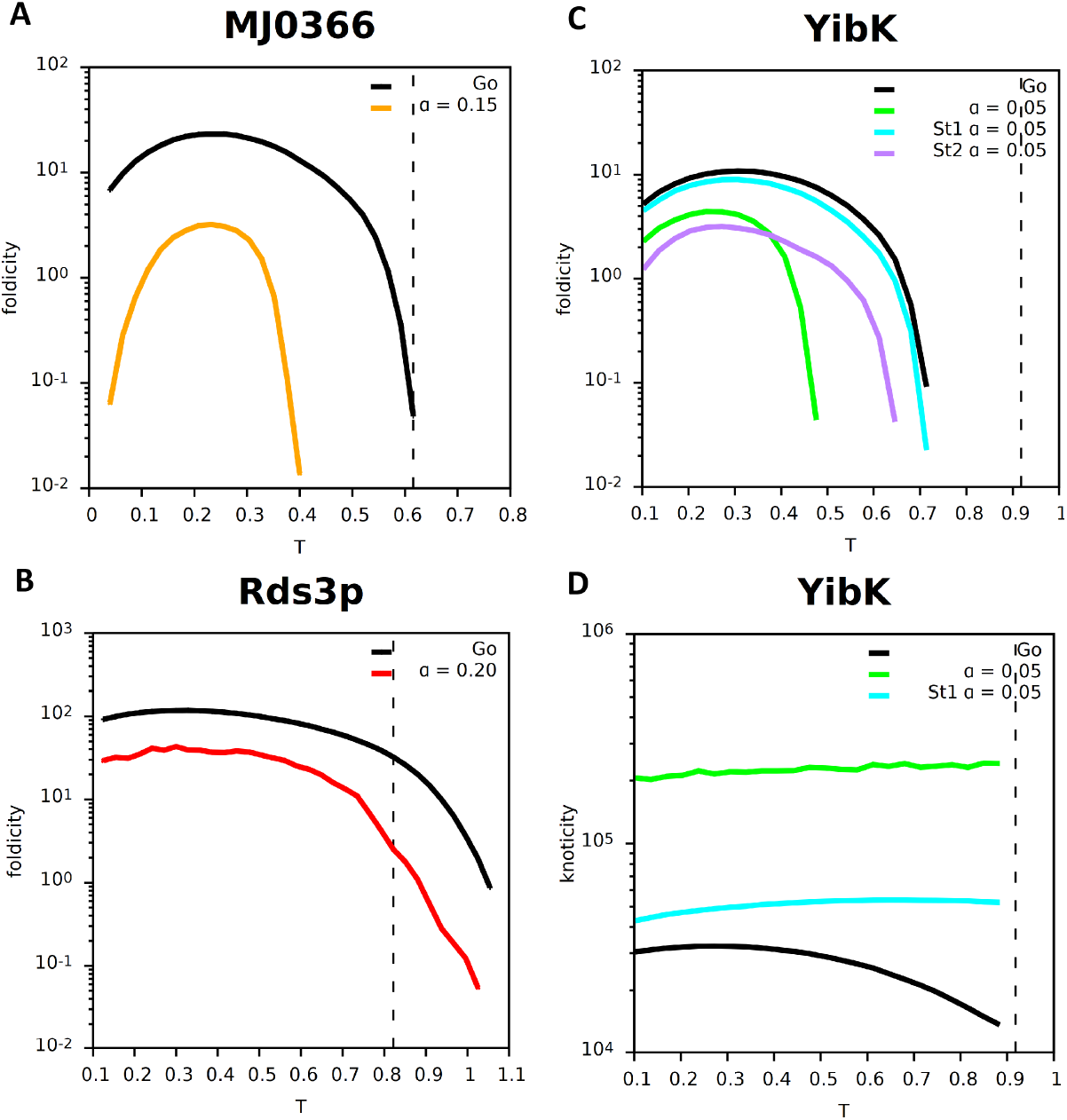
Folding efficiency. Dependence of the foldicity on temperature for proteins MJ0366 (A), Rds3p (B) and YibK (C). Also shown is the dependence of knoticity on temperature for protein YibK (D). In the case of YibK, St1 refers to the modified variant of the Gō potential that considers specific non-native interactions only between stretches (75-87) and (119-122), while St2 considers only the stretches (86-93) and (122-147). The vertical dashed line indicates the melting temperature, *T*_*m*_, for the Gō potential.

## 4 Non-native interactions and folding efficiency

Here we explore the potential role of non-native interactions in assisting the folding transition of knotted proteins. In order to do so, it is necessary to conserve the linear topology of the chain by means of MC simulations at fixed temperature. We will assess the role of non-native interactions in assisting the formation of a fully folded conformation and, in the case of protein YibK, in assisting the formation of a knotted conformation (as explained in the Methods section, in this case we are interested in the transition to the first knotted conformation independently of its degree of structural consolidation). Structural consolidation is assessed according to the criteria in the Methods section, a conformation being accepted as unfolded if it is unknotted and its *Q* is smaller than 0.25 for MJ0366, 0.26 for Rds3p and 0.34 for YibK, and being accepted as folded if it is knotted and its *Q* is greater than 0.58 for MJ0366, 0.61 for Rds3p and 0.72 for YibK. To carry out these tasks we define *foldicity* as being the number of folding transitions occurring per million mcs (Mmcs) at each considered temperature, and *knoticity* as being the number of knotting transitions occurring per million mcs (Mmcs) at each considered temperature. Apart from considering the effects of non-native interactions added in a sequence-specific manner, in the case of protein YibK, we will also consider the effect of specific non-native interactions, namely, those that have been identified in the previous section as stabilizers of knotted conformations, and those that have been identified by Shakhnovich et al. [10] as being critical to achieve 100% folding efficiency. Since the addition of non-native interactions in a sequence-dependent manner can change the nature of the folding transition from two-state to downhill (Figure 3), we will consider the highest value of the parameter *α* (which controls the strength of non-native interactions) that conserves the two-state nature of the transition.

The results reported in Figure7 (A-C) show that the three model systems are able to fold under a pure native-centric Gō potential, but this is only observed below a certain threshold temperature which is well below the melting temperature, *T*_*m*_, in the case of protein YibK. Furthermore, there is an optimal temperature at which folding proceeds at maximum efficiency. Thus, in the context of the presently adopted model and sampling protocol, non-native interactions are not necessary to efficiently fold a knotted protein, provided the temperature range of the simulations is adequate.

The results reported in Figure7 (A-C) also show the somehow surprising result that the deep knot in protein Rds3p does not hamper its folding efficiency, which is actually higher than that of protein MJ0366 (featuring a shallow knot) across the considered temperature range. It was suggested, based on results obtained in the scope of lattice models, that local structure (i.e. order) inside the hydrophobic core strongly inhibits entanglements and knots [42]. The better folding efficiency of Rds3p may thus be explained by the fact that it features a lower secondary structural content in the core in comparison to MJ0366.

On the other hand, the addition of sequence-specific non-native interactions clearly hampers folding, with foldicity decreasing up to one order of magnitude in the considered model systems. This may be due to the fact that non-native interactions persist in nearly native (i.e. stable) conformations thus precluding the formation of native ones; but the latter must be established in order for the adopted folding criterion to be satisfied. Consequently, more MC steps are required for a simulation to finish which reduces the total number of folding transitions observed per Mmcs. Focusing on the results obtained for YibK, we note that the non-native interactions establishing between the two stretches, (75-87) and (119-122), of the polypeptide chain, do not significantly alter the knotting efficiency relative to the native centric potential. However, foldicity decreases for the specific non-native interactions considered in [10]. This likely resides in specificities of the model and simulation protocol adopted in that case.

A completely different scenario is observed when one looks into the effects of non-native interactions on knoticity of YibK. In this case, the addition of non-native interactions contributes to increase the knotting efficiency and the latter is substantial (about one order of magnitude higher across the considered temperature range) for the sequence-dependent potential. We observed that on average the firstly formed knotted conformations have *Q* ≈ 0.3. The added non-native interactions are thus capable of energetically stabilizing partially folded conformations and compensate for the entropy loss due to knotting. We thus conclude that non-native interactions assist knotting but are not a pre-requisite to efficiently fold knotted proteins.

## 5 Conclusions

Determining the role of non-native interactions in folding dynamics, kinetics and mechanisms is a classical problem in protein folding, which has been extensively investigated in the scope of molecular simulations of protein models with different resolutions. It has been suggested that non-native interactions participate directly [43] or indirectly [44] in the formation of the folding nucleus, that they accelerate or decelerate the folding rate depending on whether they are more populated in the transition state or unfolded state [20, 45], that they may be determinant for protein folding cooperativity [46], that they interplay with native geometry to determine folding kinetics [47, 48], and that they determine the folding pathway at the fine level of contact formation [49].

More recently, this problem has witnessed a renewed interest in light of the hypothesis that knotted proteins require them to fold efficiently. Determining the role of native and non-native interactions for the spontaneous folding of knotted proteins has been pointed out as an important fundamental question in the literature [8].

Here, we explored the role of non-native interactions in the thermodynamics and kinetics of the folding transition of three knotted proteins featuring a trefoil knot in their native structure. In one protein, MJ0366, the knot is shallow, for the other two proteins, Rds3p and YibK, the knot is deep. We observed that non-native interactions can lower the free energy barrier to folding as observed for some unknotted proteins. Results from equilibrium simulations further indicate that non-native interactions energetically stabililise knotted conformations in a particular way: They are mainly non-local interactions being located in the vicinity of native interactions. This observation suggests that non-native interactions operate as a system of redundant interactions that assist the knotting step. In line with this finding, the (non-equilibirum) kinetic simulations of deeply knotted protein YibK show that knotting efficiency, measured as the number of knotting transitions per million MC steps, increases substantially in the presence of sequence-specific non-native interactions. The knotted conformations are partially folded conformations with an average *Q* ≈ 0.3. These results are consistent with those reported in [12] for protein AOTCase, featuring a deep trefoil knot, in which a sequence-dependent interaction potential was shown to substantially increase the knotting frequency in partially folded conformations, attaining a maximum value in conformations with *Q* ≈ 0.3.

The results from the kinetic simulations also show that, in the context of the present model and simulation protocol, native interactions alone can efficiently fold a knotted protein, provided the simulation temperature is below (or well below) the equilibrium melting temperature (for protein YibK). This is a critical finding of the present study, which may be invoked to rationalize the lack of folding success in studies of the same protein carried out by others [10, 11]. They also suggest that folding a protein with a deep knot is not necessarily less efficient than folding a protein with a shallow knot. Indeed, protein Rds3p folds more efficiently than MJ0366, presumably as a consequence of having an hydrophobic core exhibiting a lower degree of local structural order (i.e. featuring less secondary structural content) [42]. Overall, our results indicate that a sequence-specific interaction that accounts for non-native interactions may facilitate the transition from unknotted to knotted in partially folded conformations, but it may not facilitate the transition to the final folded and knotted conformation. This is likely a consequence of persistent non-native interactions in nearly folded conformations becoming highly stabilized in the temperature range where knotting is favored.

## Appendix

Let us consider a protein formed by N residues. Its native structure is defined by the set of residue coordinates 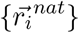, and its primary structure is defined by *{σ*_*i*_*}*, with *σ*_*i*_ being the chemical species of residue *i* = 1, …, *N*. Since there are 20 different amino acids we may assign an integer, say *l* = 1, …, 20, to each chemical species and then *σ*_*i*_ will be an integer in this range.

All pairs of non-bonded residues may potentially form a contact. Those pairs that are in contact in the native structure are native contacts. All other pairs, whenever they form a contact, are non-native contacts. We can thus determine for each pair of non-bonded residues, *i* and *j*, their chemical species *σ*_*i*_ = *l* and *σ*_*j*_ = *m* and also if they form a native or a non-native contact should they come into contact. For each pair of chemical species, *l* and *m*, we can thus count the total number of native contacts that involve this particular pair of chemical species, let ‘s call it *p*_*lm*_, and the total number of non-native contacts that also involve them, *q*_*lm*_.

In a sequence-dependent (SD) potential, the interaction energy between residues *i* and *j*, at the bottom of the potential well, *ε*_*ij*_, depends only on the chemical species, *σ*_*i*_ = *l* and *σ*_*j*_ = *m*. Let ‘s call it *I*_*lm*_. All interaction energies thus belong to a 20 *×* 20 matrix and the well bottom interaction energy of contact *i* − *j* is an element of this matrix: 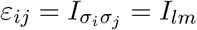.

On the other hand, the structure-based Gō potential has a well bottom contact energy of 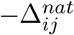. To determine the SD potential that best approximates the Gō potential we minimize the quadratic deviation

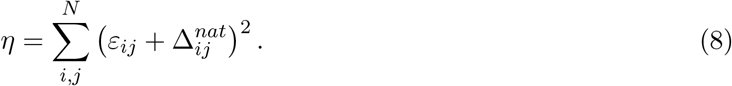

In the above sum each native contact that involves chemical species l and m contributes a term (*I*_*lm*_ + 1)^2^ and each non-native contact that involves the same chemical species a *I*_*lm*_ ^2^ term. Hence, taking into account the definitions of *p*_*lm*_ and *q*_*lm*_, the quadratic deviation sum above can be rewritten as

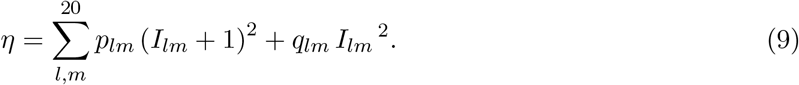

Given that the I_*lm*_ are the degrees of freedom of this minimization problem, the global minimum is achieved when

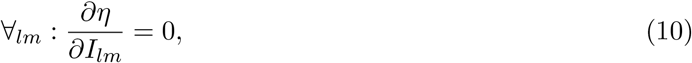

and thus, when

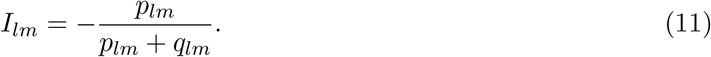

For a purely sequence-dependent (SD) potential, the total energy of a conformation defined by bead coordinates 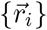 is then given by

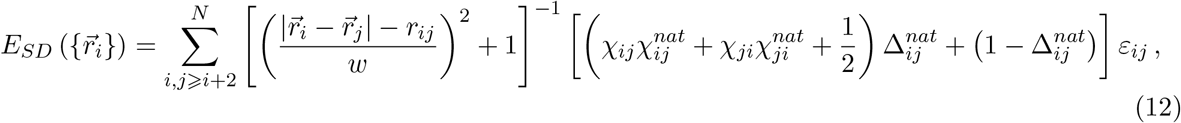

where, well bottom interaction energies 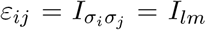 are as determined above and, if *i* − *j* is a native contact, 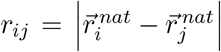 and, if *i* − *j* is a non-native contact, *r*_*ij*_ is approximated by the arithmetic mean of all native contact distances.

## 6 Author contributions

PFNF designed the research. JNCE performed the calculations. JNCE and PFNF analyzed the data. PFNF and JNCE prepared the figures and wrote the paper.

## Acknowledgments

Work supported by UID/MULTI/04046/2020 Research Unit grant from FCT, Portugal (through BioISI). JNCE acknowledges financial support from FCT, Portugal, through PhD grant SFRH/BD/144345/2019. PFNF and JNCE would like to acknowledge the contribution of the COST Action CA17139. A part of this work was performed on the computational resources of INCD (http://www.incd.pt) funded by FCT and UE under project LISBOA-01-0145-FEDER-022153. Access was granted by FCT through project 2022.26279.CPCA.A0.

## Notes

### Competing Interest Statement

The authors have declared no competing interest.

